# The Adipomyokine Follistatin-like-1 Restores Cardiovascular Function in a Swine Model of Diabetic Myocardial Infarction

**DOI:** 10.1101/2025.10.15.679868

**Authors:** Pilar Ruiz-Lozano, Shannon C. Kelly, Kleiton A. S. Silva, Taylor J. Kelty, Amira Amin, Pamela K. Thorne, Christina M. Mueller, Junqing Justin Zhang, Jan R. Ivey, Brittany M. Mezzencella, Ashley N. Clark, Boxiong Deng, Sui Wang, R. Scott Rector, Craig. A. Emter, Mark Mercola, Darla L. Tharp

## Abstract

Obesity, diabetes, and metabolic syndrome increase the incidence and complicate the management of myocardial infarction (MI). Current treatments do not adequately blunt the progression to chronic ischemic heart disease and heart failure, making these diseases the largest cause of mortality worldwide. Insulin resistance and metabolic syndrome were induced in obese Ossabaw swine prior to ischemia/reperfusion injury-induced MI. Subcutaneous administration of the adipomyokine Follistatin-like-1 (recombinant non-glycosylated human FSTL1, ngFSTL1) for two weeks, beginning one month following myocardial infarction, decreased infarct necrosis, increased blood flow, and increased cardiomyocyte proliferation within the infarct region, leading to improved systolic and diastolic function. ngFSTL-1 also enhanced coronary and peripheral vascular function by increasing BK_Ca_ channel-dependent vasodilatory capacity. We conclude that subcutaneous ngFSTL1 ameliorated clinically relevant parameters of cardiac and vascular dysfunction in a preclinical swine model of diabetic MI, and suggest FSTL1 treatment may be broadly efficacious in humans.

**ARTICLE HIGHLIGHTS:**

- Subcutaneous administration of recombinant human non-glycosylated FSTL1 (ngFSTL1) for 2 weeks, 1 month after infarction/reperfusion injury, improved cardiac function in a preclinical obese swine model of diabetic MI.
- ngFSTL1 treatment induced cardiomyocyte proliferation and improved coronary perfusion, resulting in reduced necrosis of the infarct region.
- ngFSTL1 treatment increased blood flow through peripheral arterioles, including cerebral and skeletal muscle, in addition to cardiac arterioles.
- Systemic delivery of the non-glycosylated form of the adipomyokine FSTL1 might be an effective treatment for diabetic MI.

## INTRODUCTION

The increasing prevalence of obesity and diabetes mellitus (DM) worldwide presents a growing burden on patients and healthcare systems alike. The impact of obesity, DM, and associated metabolic syndrome is largely driven by its cardiovascular complications, which include coronary artery disease, myocardial infarction, peripheral vascular disease, and neuropathy ^1^. Associated risks, such as dyslipidemia and hypertension, increase the incidence of cardiovascular disease and worsen outcomes. Consequently, diabetic patients have a worse incidence of myocardial infarction (MI), morbidity, and re-infarction relative to non-diabetics, culminating in a one-year mortality of nearly 50% after the first MI ^2-5^.

Several pathogenic mechanisms compound the effects of MI in diabetic patients. These include deranged metabolism and calcium handling that alter cardiac structure and function leading to systolic and diastolic dysfunction ^6,7^. In addition, damage to cardiac and peripheral vasculature aggravate the cardiac effects and complicate patient management ^8^.

Follistatin-like-1 (FSTL1; also known as FRP1, FSL1, OCC1 and Tsc36) is a myokine and adipokine with distant homology to follistatin, SPARC, and other FS domain proteins. FSTL1 is secreted by skeletal muscle following endurance exercise and acts on the heart to increase myocardial angiogenesis and cardiac performance ^9^. The heart also produces FSTL1 acutely after ischemic injury in the infarct region ^10^, where it promotes angiogenesis and decreases injury-related cardiomyocyte apoptosis ^11^. Glycosylation of FSTL1 can profoundly affect its function. For instance, the naturally glycosylated, circulating FSTL1 activates cardiac fibrosis ^12^, limits cardiomyocyte apoptosis ^13,14^, and might modulate mitochondrial function ^15^. In contrast, non-glycosylated or partially glycosylated forms of FSTL1 have been shown to remuscularize the infarct region when delivered directly to the heart following myocardial infarction in non-diabetic mice and pigs ^10,16^.

For this study, we evaluated whether non-glycosylated FSTL1 (ngFSTL1) could be delivered systemically, via subcutaneously implanted osmotic pump, to improve cardiac morphology and function following MI in a swine model of obesity and metabolic dysfunction ^17^. To our knowledge, this is the first study to induce ischemia/reperfusion induced injury and MI in a diabetic swine model. One month following MI, a 2 week regimen of subcutaneous ngFSTL1 delivery improved cardiac function, decreased myocardial necrosis, and increased myocardial and peripheral perfusion. Thus, subcutaneously administered ngFSTL1 in the setting of diabetic MI might be an effective strategy to restore heart function and improve central and peripheral vascular perfusion in the heart, skeletal muscle, and brain.

## RESEARCH DESIGN AND METHODS

### Animals

All animal protocols were approved by the University of Missouri Animal Care and Use Committee and were in accordance with the “Principles for the Utilization of Care of Vertebrate Animals Used in Testing Research and Training”. Intact female Ossabaw swine (CorVus Biomedical, LLC, Crawfordsville, IN, USA) were placed on a high-fat/fructose/cholesterol Western Diet (KT324, CorVus Biomedical, LLC) containing 16.3% kCal from protein, 42.9% kCal from fat, and 40.8% kCal from carbohydrates at approximately 2 months of age. Animals were fed 1 kg once per day, and water was supplied ad libitum.

### Ischemia/Reperfusion Induced Myocardial Infarction

Ischemia/reperfusion-induced myocardial infarction was performed at 6 months of age by inflation of an angioplasty balloon in the left anterior descending (LAD) coronary artery as previously described ^18,19^ with slight modifications (see Supplementary Methods). The LAD was occluded distal to the 1^st^ diagonal branch (D1) for 90 minutes, followed by reperfusion. Transthoracic echocardiography was performed at baseline, pre- and immediately post-MI, 1-month post-MI, and 2-months post-MI. Short-axis two-dimensional M-mode images were collected at the mid-papillary level of the LV and were analyzed using GE EchoPac Software.

### ngFSTL1 delivery

One month following MI, Alzet Osmotic Pumps were implanted subcutaneously on both lateral surfaces of the neck. One Alzet pump released FSTL1 (120 mg FSTL1 in 2 ml TBS, released 5.0 uL per hour over 14 days) or vehicle (TBS, Tris Buffered Saline). The other Alzet pump released EdU (12 mg EdU in 2 ml PBS, released 2.5 uL per hour for 28 days) to label proliferating cells or vehicle (PBS, Phosphate Buffered Saline). See Supplementary Methods for details.

### Assessment of cardiac and vascular physiology

Pressure-Volume (PV) loops, regional blood flow, and pressure myography were performed as described in the Supplementary Methods section. Mitochondrial function was assessed in isolated mitochondria as previously described ^20^. Mitochondrial respiration was normalized to protein concentration obtained from BCA assay per manufacturer’s instructions. Investigators performing all physiological analyses were blinded with respect to cohort identity.

### Statistical Analyses

All data were graphed and analyzed using GraphPad Prism 10. Pressure myography data, mitochondrial data, and serial echocardiography data were analyzed using repeated measures ANOVA; all other two-group comparisons (MI vs MI + FSTL1) were analyzed using unpaired t-tests (two-tailed). Outliers were determined using ROUT with Q=5%. Data were considered significant at ^*^p<0.05 and †p<0.1.

### Data and Resource Availability

Data and resources are available on request from the corresponding authors.

## RESULTS

### Porcine model of ischemia/reperfusion-induced myocardial infarction leading to heart failure with reduced ejection fraction (HFrEF)

Obesity, metabolic syndrome, insulin resistance, and hypercholesterolemia were induced by placing Ossabaw swine on a high fat/fructose/cholesterol Western Diet for 4 months (**Figure 1a)** and was confirmed by multiple parameters (as previously reported by ^17^), including insulin resistance with increased HOMA-IR (**Figure 1d**, 3.24 ± 0.24 compared to 0.76 ± 0.07 for lean control non-MI Ossabaws; p<0.05). Myocardial infarction was induced by inflating a balloon catheter distal to the first diagonal branch of the left anterior descending (LAD) coronary artery for 90 minutes, followed by balloon deflation to mimic ischemia/reperfusion (**Figure 1b,c**). Western diet was initiated 4 months prior to myocardial infarction and continued for the duration of the study (2 months post-MI).

**Figure 1.**
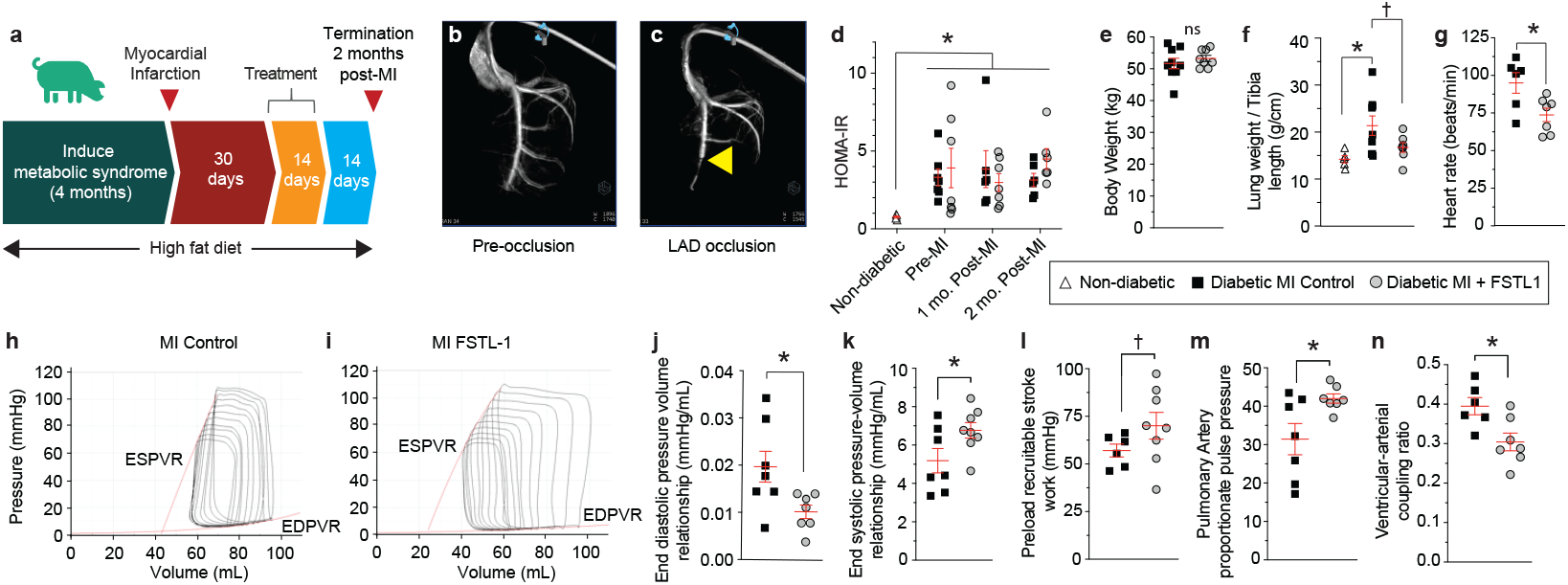
Subcutaneous ngFSTL1-treatment of diabetic MI reduced pulmonary edema, prevented diastolic dysfunction, and improved contractility. **a)** Schematic of intervention. **b,c)** Representative angiogram before and after balloon occlusion. Yellow arrowhead indicates point of occlusion. **d)** HOMA-IR measurements at time of MI and 1 and 2 months post-MI. **e-g)** morphometric measurements of body weight, wet lung weight, and heart rate 2 months post-MI. **h,i)** Representative PV Loops. **j-m)** ngFSTL1 treatment decreased EDPVR (j), increased ESPVR (k) and PRSW (l), reduced the narrowing of pulse pressure, calculated as pulmonary artery pulse pressure/pulmonary systolic pressure, and graphed as % (m), and improved ventricular-arterial coupling (n). †, p<0.01; ^*^, p<0.05; unpaired t-test.

Four weeks after MI, the ischemia/reperfusion cohort was randomized into control (vehicle; n=9) and treatment (FSTL1; n=8). The ejection fraction in the MI group at this timepoint was 39.9 ± 1.8%, which was significantly less than that in non-MI control pigs (51.92 ± 3.1%; p<0.05). The four week lapse between MI and treatment initiation was selected to confirm decay of cardiac function in response to MI and evaluate the potential recovery following treatment.

### Subcutaneous delivery of ngFSTL1 after MI improved cardiac function and prevented wall thinning

The I/R plus treatment cohort received 8.5 mg/day ngFSTL1 infused from subcutaneously implanted Alzet osmotic pumps for 14 days while the I/R no treatment cohort received pumps with PBS vehicle. The treatment did not affect insulin resistance as HOMA-IR levels remained elevated throughout the study (**Figure 1d**). At termination, vehicle-treated Ossabaw pigs presented increased wet lung weight (p<0.05), a hallmark of heart failure, which was decreased by the ngFSTL1 treatment (p<0.1; **Figure 1e-f)**. There were no changes in body surface area, tibia length, or heart weight (**Supplementary Figure S1**).

In-depth analysis of left ventricular hemodynamics by pressure-volume (PV) loops across different loads (representative PV Loops shown in **Figure 1h,i**) demonstrated that ngFSTL1 significantly reduced end-diastolic pressure volume relationship (EDPVR, **Figure 1j**), with no change in end systolic nor end diastolic pressures or volumes at rest (**Supplementary Figure S2a-d**). A reduction in EDPVR suggests ngFSTL1 prevents diastolic dysfunction, which occurs more commonly in diabetic than non-diabetic patients after MI and is associated with worse outcomes ^21^.

ngFSTL1 improved contractile function, as evidenced by increased end systolic pressure-volume relationship systolic function (ESPVR, **Figure 1k**) and an increased preload recruitable stroke work (PRSW, **Figure 1l**). We did not observe a significant increase in EF% or stroke volume after the two week treatment (**Supplementary Figure S2e,f**). Echocardiography, however, revealed reduced left ventricular internal end systolic and diastolic dimensions **(Supplementary Figure S3a,b)**, consistent with the improved contractility observed by PV hemodynamics. Further, ngFSTL1 had a normalizing effect on pulmonary artery proportionate pulse pressure (PAPPP) (**Figure 1m**) and on ventricular-arterial coupling ratio (**Figure 1n**), indicative of improved ventricular-vascular interactions.

### ngFSTL1 reduced myocardial necrosis and adverse remodeling after MI

ngFSTL1 treatment blunted the decrease in LV systolic septal wall thickness relative to control diabetic MI pigs **(Supplementary Figure S3c)**, suggesting improved contractility and myocardial perfermance as a possible mechanism for functional recovery. Examination of gross tissue slices (**Figure 2a,b**) revealed that, although the initial infarct region was comparable across cohorts (16 ± 4% of the LV volume in controls versus 15 ± 4% in treated animals, (**Figure 2c**), there was a marked decrease in the extent of necrosis in the infarct region of ngFSTL1 treated animals relative to control MI hearts (**Figure 2d**) as well as decreased iron deposition (**Figure 2e-g**). Together, these data indicate that ngFSTL1 blunts or reverses adverse remodelling.

**Figure 2.**
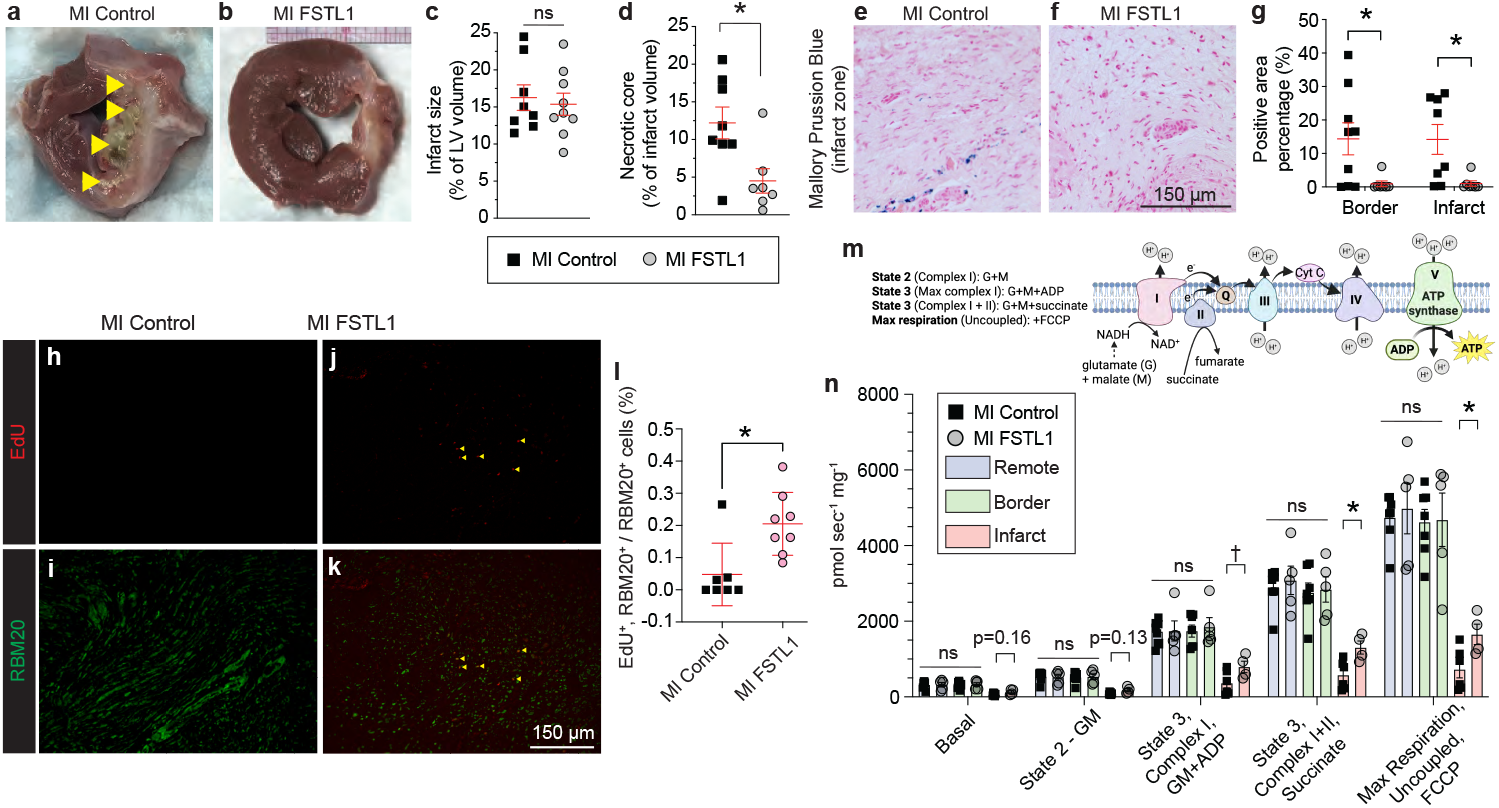
Subcutaneous ngFSTL1 treatment of diabetic MI restored heart morphology and mitochondrial function. **a,b)** Representative images of infarct. Yellow arrowheads denote necrotic core. **c,d)** ngFSTL1-treated diabetic animals displayed reduced necrotic core area within the infarct. **e-g)** ngFSTL1-treated animals displayed decreased iron (Mallory Prussian Blue staining) in both border and infarct areas. **h-l)** ngFSTL1 treatment stimulated EdU incorporation in cardiomyocytes, as determined by co-staining of EdU (red, h,j) with the cardiomyocytes specific marker RBM20 (green i,k) and as quantified (l) relative to total RBM20. **m)** Schematic of mitochondrial electron transport chain complexes assayed. **n)** Quantification of the cardiac mitochondrial respiration in the infarct zone, which is increased in ngFSTL1-treated diabetic pigs selectively within the infarct region. †, p<0.01; ^*^, p<0.05; unpaired t-test.

Previous studies had shown that ngFSTL1 can induce cardiomyocyte proliferation in infarcted myocardium in non-diabetic animals ^10,16^. To test for evidence of cardiomyocyte proliferation, we administered the nucleotide analogue EdU simultaneously with ngFSTL1 or vehicle control from a separate Alzet pump (see Supplementary Methods). EdU incorporation into cardiomyocytes (unambiguously identified based on RBM20 immunostaining) was quantified in histological sections 2 months after MI. ngFSTL1-treated animals showed a significantly higher prevalence of EdU^+^ cardiomyocytes than did untreated cohorts (**Figure 2h-l**), suggesting ngFSTL1 promotes cardiomyoycyte proliferation in the diabetic MI setting. EdU incorporation in remote myocardium was not observed (not shown but as previously reported ^10^).

Consistent with a localized effect in the infarct region, mitochondrial respiratory activity was increased within the infarct zone (**Figure 2m,n**). ngFSLT1 increased State 3, Complex I – GM + ADP (non-significant), State 3, Complex I + II – Succinate (^*^p<0.05, RM ANOVA state x group interaction), and maximal uncoupled (FCCP;^*^p<0.05, unpaired t-test) respiration, thus improving complex 2-dependent ADP stimulated mitochondrial respiration, consistent with the elevated ATP demains of improved systolic function. No changes were observed in the remote or border zones (**Figure 2n**), consistent with a localized effect of ngFSTL1 despite systemic delivery.

### Peripheral vascular effects

ngFSTL1-treatment improved myocardial blood flow, as demonstrated by samarium-labeled microsphere measurements. Coronary blood flow was increased throughout the LV, but not the mid-septum or remote RV (**Figure 3a** and **Supplementary Figure S4**). ngFSTL1 similarly increased blood flow in skeletal muscle and brain (**Figure 3b,c** and **Supplementary Figure S4**). To establish a mechanism for the increased blood flow, we performed ex vivo measurements of arteriole constriction and dilation (**Figure 3d,e)**. We probed the activity of large-conductance calcium-activated potassium (BK_Ca_) channels since they counter vasoconstriction ^22,23^ and are impaired in both coronary and peripheral vasculature of type 2 diabetic patients ^24,25^. Coronary arterioles isolated from the remote region of the left ventricle of ngFSTL1 animals exhibited a more potent response to NS1619, a small molecule activator of BK_Ca_ channels (**Figure 3d**). We also examined second order (2a) pial arterioles of the middle cerebral artery. ngFSTL treatment did not significantly alter the vasodilatory response to NS1619 alone nor affect U46619 (thromboxane A2 receptor agonist)-mediated constriction (**Supplementary Figure S5a,b**). However, ngFSTL1 significantly enhanced the ability of NS1619 to limit U46619-mediated constriction (**Figure 3e; Supplementary Figure S5c,d)**. These data suggest that one of the mechanisms by which systemic ngFSTL1 increases peripheral blood flow is by enhancing BK_Ca_ channel activity to counteract vasoconstriction (**Figure 3f)**.

**Figure 3.**
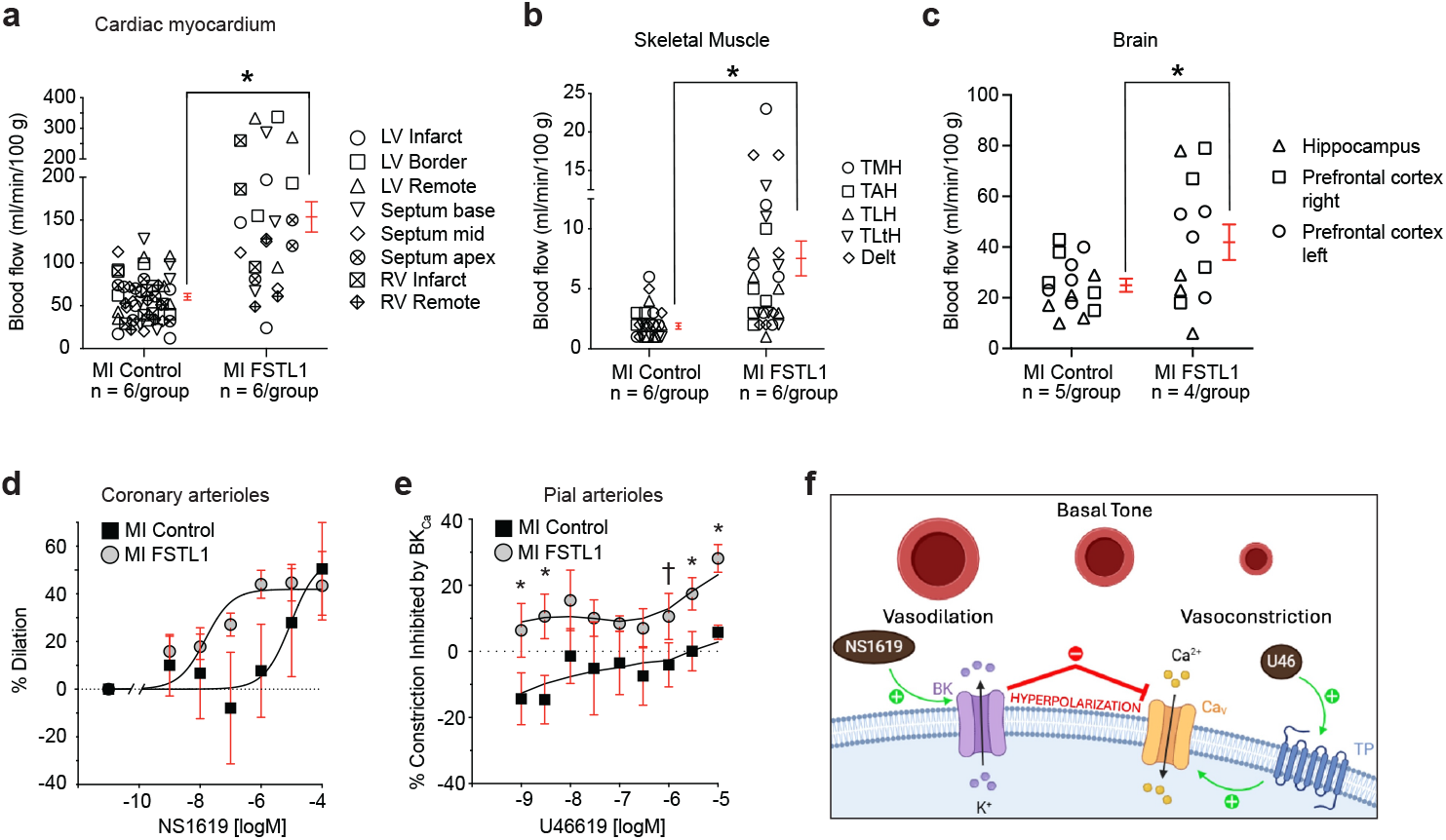
Subcutaneous treatment of diabetic MI with ngFSTL1 improved coronary and peripheral vascular function. **a,b)** ngFSTL1-treatment improved blood flow in coronary (a), skeletal muscle (b), and brain (c) vasculature, as measured by microsphere deposition (see Methods). Abbreviations: TMH, triceps brachii medial head; TAH, triceps brachii anterior head; TLH, triceps brachii long head; TLtH, triceps brachii lateral head; Delt, deltoid. **d)** Coronary arterioles isolated from ngFSTL1-treated hearts showed increased dilatory response to NS1619, a BK_Ca_ channel activator. **e)** Second order (2a) pial arterioles of the middle cerebral artery from ngFSTL1-treated animals showed a heightened dilatory response to the BK_Ca_ activator NS1619 that opposed vasoconstriction induced by the thromboxane A2 agonist U46619 (graphed as % of U46619 constriction inhibited by NS1619). **f)** Together, these data indicate that ngFSTL1 promotes vasodilation through activation of BK_Ca_ channels. †, p<0.01; ^*^, p<0.05; unpaired t-test (a-c), RM ANOVA main effect of group (p<0.05 d,e).

## DISCUSSION

In this study, we used Ossabaw swine, which are genetically predisposed to develop obesity, insulin resistance, and metabolic syndrome, as a preclinical model of diabetic MI. MI in diabetic patients is associated with an enhanced risk of complications and mortality relative to non-diabetic patients, and acute MI continues to be a major cause of death among diabetic patients ^26^. To our knowledge, this is the first instance of modeling ischemia-reperfusion induced myocardial infarction in diabetic swine, which enhances clinical translatability relative to rodent models. By one month after initiation of treatment, subcutaneously delivered ngFSTL1 improved LV diastolic function, healed the infarct, improved ATP production, and increased myocardial blood flow, resulting in improved ventricular-vascular coupling, improved LV contractility, and normalized pulmonary pulse pressure post-MI. Notably, these effects were elicited without a change in body mass or insulin resistance, indicating direct cardiovascular effects. ngFSTL1 also increased blood flow to the myocardium as well as other peripheral vascular beds, namely skeletal muscle and brain. This is the first demonstration of the therapeutic potential of ngFSTL1 in a preclinical large animal model of diabetic MI.

At a morphological level, ngFSTL1 treatment decreased necrotic tissue volume, decreased iron overload, and promoted cardiomyocyte incorporation of EdU. It also increased mitochondrial function within the infarct zone, illustrating the selective action of systemically-administered ngFSTL1 on injured versus healthy (remote) myocardium. These results, plus the confirmation that heart function had declined at the initiation of treatment, are consistent with a restorative rather than cardioprotective mechanism of action.

In coronary arterioles, ngFSTL1 treatment enhanced the potency of the BK_Ca_ channel activator NS1619. In pial arterioles of the middle cerebral artery, ngFSTL1 did not significantly affect NS1619 alone, but potentiated its ability to counter constriction induced by the Thromboxane A2 agonist U46619. Vascular smooth muscle BK_Ca_ channels function as a brake to limit overconstriction, and loss of BKCa channel expression and activity occurs in diabetes, contributing to vascular complications, including nephropathy, coronary artery disease, and retinopathy ^24^. Activation of BK_Ca_ channel activity by systemic delivery of ngFSTL1 might alleviate vascular dysfunction in this setting. Additional studies are needed to distinguish whether ngFSTL1 acts directly on vasculature to increase BK_Ca_ activity, or elicits its effect as a consequence of enhanced cardiac function (i.e., perfusion and shear stress).

Finally, this study illustrated the efficacy of systemic administration rather than direct delivery to the heart either as non-glycosylated protein ^10^ or as modified RNA encoding an engineered form lacking N-linked glycosylation sites ^16^ as done previously. Systemic administration should enhance clinical translatability relative to direct intramyocardial delivery.

In summary, we created a novel Ossabaw swine model of diabetic MI to evaluate treatment with ngFSTL1. Subcutaneously delivered ngFSTL1 ameliorated systolic and diastolic dysfunction and improved coronary and peripheral blood flow. Previous studies had shown that direct delivery ngFSTL1 to the heart of non-diabetic animals reduced scar volume and enhanced cardiac contractility following MI ^10,16^. This present study is the first to show the therapeutic efficacy of systemically delivered ngFSTL1 and the first to show efficacy in diabetic MI. Treatment was initiated one month after MI, suggesting that ngFSTL1 might be an effective and straightforward therapeutic to treat cardiac and vascular complications of diabetic MI.

## Acknowledgements

The University of Missouri Veterinary Medical Diagnostic Lab performed histology. This work was also supported in part with resources and equipment from the Truman VA Memorial Hospital.

## Author contributions

PRL: Conception and design, acquisition of funding, analysis and interpretation of data, writing and editing the manuscript.

PKT: Acquistion, analysis, and interpretation of data

CMM: Acquisition and analysis of data

JJZ: Acquisition and analysis of data

JRI: Acquisition and analysis of data

SCK: Acquisition and analysis of data

KASS: Acquisition and analysis of data

BD: Acquisition and analysis of data

AA: Acquisition and analysis of data

BMM: Acquisition and analysis of data

ANC: Acquisition and analysis of data

TJK: Acquistion, analysis, and interpretation of data, and editing of manuscript

SW: Acquisition and interpretation of data.

RSR: Acquistion, analysis, and interpretation of data, and editing of manuscript

CAE: Conception and design, administration, analysis and interpretation of data, and editing the manuscript.

MM: Conception and design, acquisition of funding, analysis and interpretation of data, writing and editing the manuscript.

DLT: Conception and design, administration, analysis and interpretation of data, writing and editing the manuscript.

## References

1 Lowenstern, A. et al. Influence of Obesity on Coronary Artery Disease and Clinical Outcomes in the ADVANCE Registry. Circ Cardiovasc Imaging 16, e014850, doi:10.1161/CIRCIMAGING.122.014850 (2023).

2 Matsushita, K. et al. Prognostic impact of diabetes mellitus on in-hospital mortality in patients with acute myocardial infarction complicating renal dysfunction according to age and sex. Hellenic J Cardiol, doi:10.1016/j.hjc.2023.11.002 (2023).

3 Blin, P. et al. Patients with stable coronary artery disease and type 2 diabetes but without prior myocardial infarction or stroke and THEMIS-like patients: real-world prevalence and risk of major outcomes from the SNDS French nationwide claims database. Cardiovasc Diabetol 20, 229, doi:10.1186/s12933-021-01416-1 (2021).

4 Awad, H. H. et al. Magnitude, treatment, and impact of diabetes mellitus in patients hospitalized with non-ST segment elevation myocardial infarction: A community-based study. Diab Vasc Dis Res 13, 13–20, doi:10.1177/1479164115609027 (2016).

5 Colombo, M. G. et al. Association of obesity and long-term mortality in patients with acute myocardial infarction with and without diabetes mellitus: results from the MONICA/KORA myocardial infarction registry. Cardiovasc Diabetol 14, 24, doi:10.1186/s12933-015-0189-0 (2015).

6 El Hadi, H., Vettor, R. & Rossato, M. Cardiomyocyte mitochondrial dysfunction in diabetes and its contribution in cardiac arrhythmogenesis. Mitochondrion 46, 6–14, doi:10.1016/j.mito.2019.03.005 (2019).

7 Hamilton, S. & Terentyev, D. Proarrhythmic Remodeling of Calcium Homeostasis in Cardiac Disease; Implications for Diabetes and Obesity. Front Physiol 9, 1517, doi:10.3389/fphys.2018.01517 (2018).

8 Russo, M. P., Grande-Ratti, M. F., Burgos, M. A., Molaro, A. A. & Bonella, M. B. Prevalence of diabetes, epidemiological characteristics and vascular complications. Arch Cardiol Mex 93, 30–36, doi:10.24875/ACM.21000410 (2023).

9 Xi, Y., Gong, D. W. & Tian, Z. FSTL1 as a Potential Mediator of Exercise-Induced Cardioprotection in Post-Myocardial Infarction Rats. Sci Rep 6, 32424, doi:10.1038/srep32424 (2016).

10 Wei, K. et al. Epicardial FSTL1 reconstitution regenerates the adult mammalian heart. Nature 525, 479–485, doi:10.1038/nature15372 (2015).

11 Ouchi, N. et al. DIP2A functions as a FSTL1 receptor. J Biol Chem 285, 7127–7134, doi:10.1074/jbc.M109.069468 (2010).

12 Maruyama, S. et al. Follistatin-like 1 promotes cardiac fibroblast activation and protects the heart from rupture. EMBO Mol Med 8, 949–966, doi:10.15252/emmm.201506151 (2016).

13 Ouchi, N. et al. Follistatin-like 1, a secreted muscle protein, promotes endothelial cell function and revascularization in ischemic tissue through a nitric-oxide synthase-dependent mechanism. J Biol Chem 283, 32802–32811, doi:10.1074/jbc.M803440200 (2008).

14 Shimano, M. et al. Cardiac myocyte follistatin-like 1 functions to attenuate hypertrophy following pressure overload. Proc Natl Acad Sci U S A 108, E899–906, doi:10.1073/pnas.1108559108 (2011).

15 Seki, M. et al. Acute and Chronic Increases of Circulating FSTL1 Normalize Energy Substrate Metabolism in Pacing-Induced Heart Failure. Circ Heart Fail 11, e004486, doi:10.1161/CIRCHEARTFAILURE.117.004486 (2018).

16 Magadum, A. et al. Ablation of a Single N-Glycosylation Site in Human FSTL 1 Induces Cardiomyocyte Proliferation and Cardiac Regeneration. Mol Ther Nucleic Acids 13, 133–143, doi:10.1016/j.omtn.2018.08.021 (2018).

17 Olver, T. D. et al. Western Diet-Fed, Aortic-Banded Ossabaw Swine: A Preclinical Model of Cardio-Metabolic Heart Failure. JACC Basic Transl Sci 4, 404–421, doi:10.1016/j.jacbts.2019.02.004 (2019).

18 Zaid, M. et al. Mechanism-Driven Modeling to Aid Non-invasive Monitoring of Cardiac Function via Ballistocardiography. Front Med Technol 4, 788264, doi:10.3389/fmedt.2022.788264 (2022).

19 Ishikawa, K. et al. Characterizing preclinical models of ischemic heart failure: differences between LAD and LCx infarctions. Am J Physiol Heart Circ Physiol 307, H1478–1486, doi:10.1152/ajpheart.00797.2013 (2014).

20 Sheldon, R. D. et al. eNOS deletion impairs mitochondrial quality control and exacerbates Western diet-induced NASH. Am J Physiol Endocrinol Metab 317, E605–E616, doi:10.1152/ajpendo.00096.2019 (2019).

21 Aronson, D. et al. Impact of diastolic dysfunction on the development of heart failure in diabetic patients after acute myocardial infarction. Circ Heart Fail 3, 125–131, doi:10.1161/CIRCHEARTFAILURE.109.877340 (2010).

22 Carlton-Carew, S. R. E. et al. Stimulation of the calcium-sensing receptor induces relaxations of rat mesenteric arteries by endothelium-dependent and -independent pathways via BK(Ca) and K(ATP) channels. Physiol Rep 12, e15926, doi:10.14814/phy2.15926 (2024).

23 Au, A. L. et al. Activation of iberiotoxin-sensitive, Ca2+-activated K+ channels of porcine isolated left anterior descending coronary artery by diosgenin. Eur J Pharmacol 502, 123–133, doi:10.1016/j.ejphar.2004.08.045 (2004).

24 Nieves-Cintron, M. et al. Impaired BK(Ca) channel function in native vascular smooth muscle from humans with type 2 diabetes. Sci Rep 7, 14058, doi:10.1038/s41598-017-14565-9 (2017).

25 Borbouse, L. et al. Impaired function of coronary BK(Ca) channels in metabolic syndrome. Am J Physiol Heart Circ Physiol 297, H1629–1637, doi:10.1152/ajpheart.00466.2009 (2009).

26 Jacoby, R. M. & Nesto, R. W. Acute myocardial infarction in the diabetic patient: pathophysiology, clinical course and prognosis. J Am Coll Cardiol 20, 736–744, doi:10.1016/0735-1097(92)90033-j (1992).

