## Supplementary Figures for "The Adipomyokine Follistatin-like-1 Restores Cardiovascular Function in a Swine Model of Diabetic Myocardial Infarction"

Supplementary Figure S1

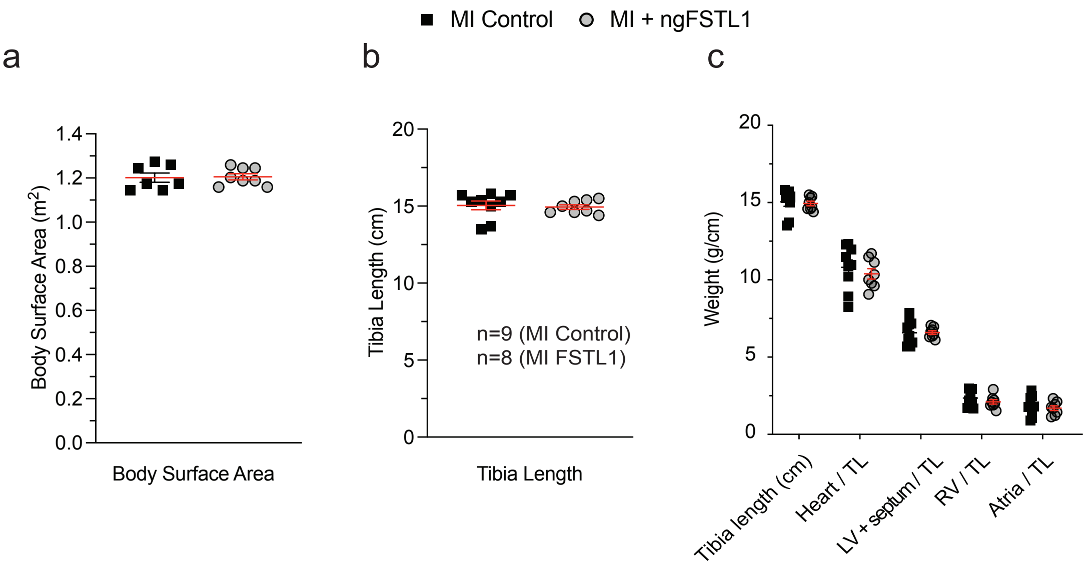


**Supplementary Figure S1. ngFSTL1 had no effect on body surface area, tibia length, nor heart weight. a-c)** MI control and MI FSTL1 animals had the same body surface area (a), tibia length (b), total heart weight, left ventricular (LV) + septum, right ventricle (RV) and atria (c) weights normalized to tibia length. See main Figure 1 for remainder of morphometric parameters.

Supplementary Figure S2

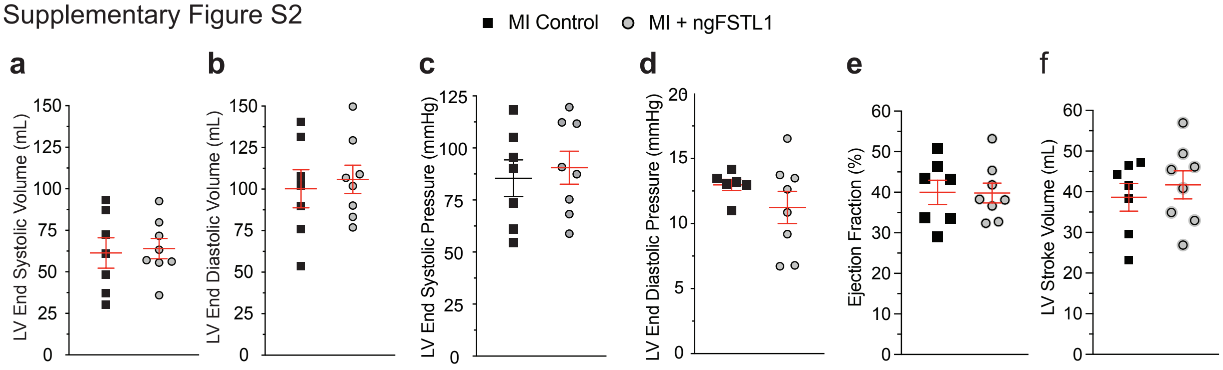


**Supplementary Figure S2. Treatment effects on hemodynamic parameters at two months after MI**. LV end systolic volume and pressure (a, c), end diastolic volume and pressure (b, d), ejection fraction % (e) and stroke volume (f) were not different between groups . Data plotted as mean ± s.e.m..

Supplementary Figure S3

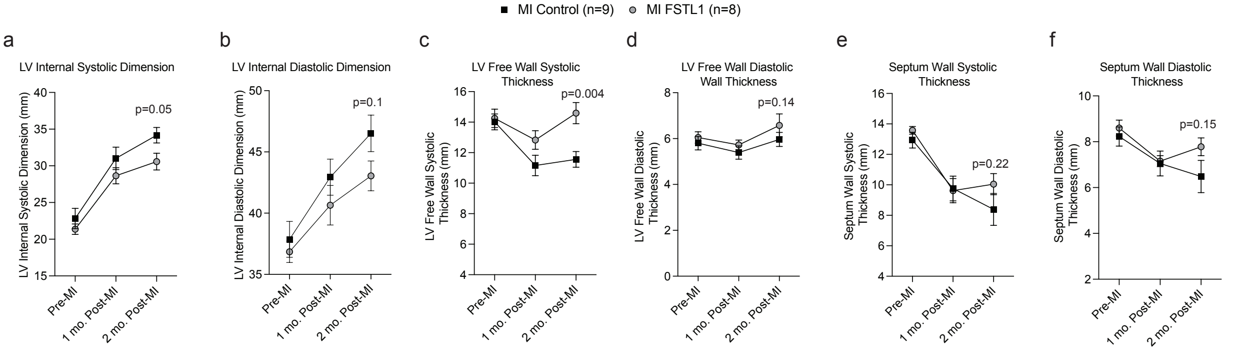


**Supplementary Figure S3. Short-axis 2D M-mode echocardiographic parameters at one- and two-months post-MI.** LV internal systolic (a) and diastolic (b) dimensions decreased with ngFSTL1 treatment. LV free wall systolic (c) and diastolic (d) as well as septal systolic (e) and diastolic (f) thicknesses increased with treatment. Data plotted as mean ± s.e.m. p-values (RM ANOVA, time x group interaction) are indicated for 2-month time points.

Supplementary Figure S4

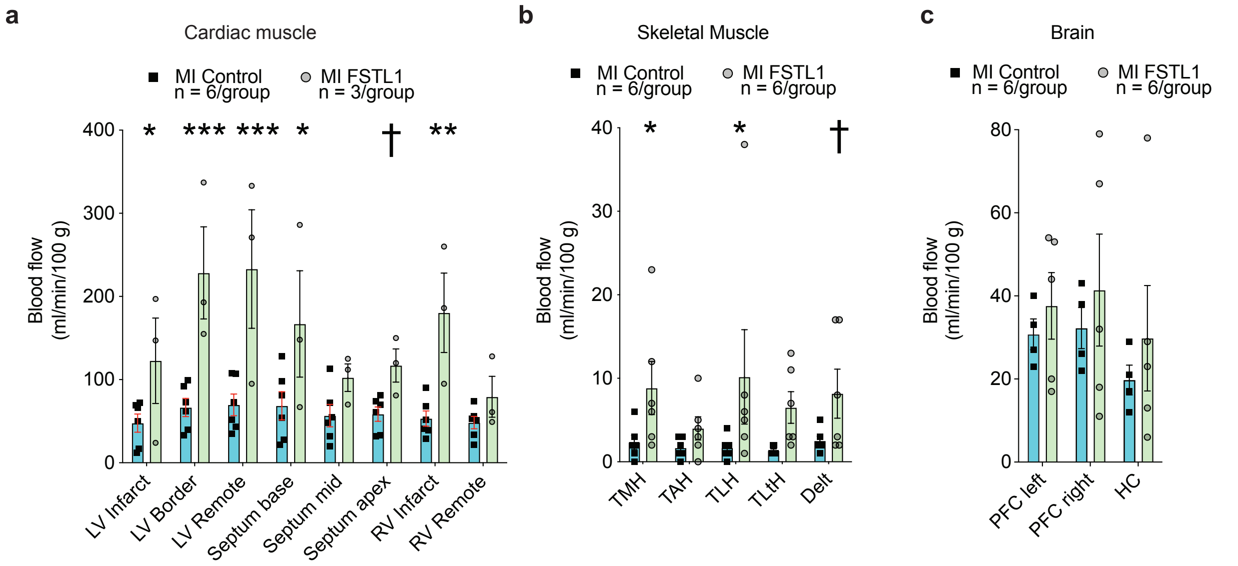


**Supplementary Figure S4. Subcutaneous treatment of ngFSTL1 in diabetic pigs improves blood flow in different tissues.** a-c) ngFSTL1 effects on blood flow in situ were measured by by samarium-labeled microsphere measurements in myocardium (a), skeletal muscle (b), and brain (c). See Methods for experimental details. Figure 3 shows aggregate data for the individual tissues. Abbreviations: PFC, Prefrontal cortex; TMH, triceps brachii medial head; TAH, triceps brachii anterior head; TLH, triceps brachii long head; TLtH, triceps brachii lateral head; Delt, deltoid. Data plotted as mean ± s.e.m. p adjusted values (ANOVA with Benjamini-Hochlberg post-hoc test for multiple comparisons) indicated by †, $p<0.1$; *, $p<0.05$; **, <0.01; ***, <0.001.

Supplementary Figure S5

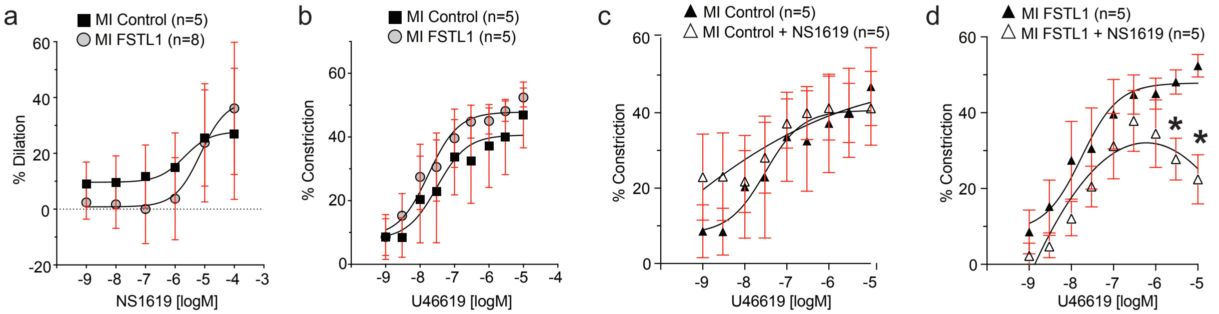


**Supplementary Figure S5. Middle cerebral artery second arteriole function. a,b)** ngFSTL1 treatment did not significantly alter the vasodilatory response to the $BK_{Ca}$ channel activator NS1619 (a) nor the vasoconstrictive response to the thromboxane A2 receptor agonist U46619 (b). **c,d)** However, ngFSTL1 treatment (MI FSTL1) potentiated the effectiveness of NS1619 to limit the vasoconstrictive activity of U46619 (d). This activity was not seen in vessels from untreated (MI Control) animals (c). Data plotted as mean ± s.e.m. *, $p<0.05$ (RM ANOVA, main effect of group).
