## Supplementary Methods for "The Adipomyokine Follistatin-like-1 Restores Cardiovascular Function in a Swine Model of Diabetic Myocardial Infarction"

**Animals.** Intact female Ossabaw swine were obtained from CorVus Biomedical, LLC in Crawfordsville, IN, USA and placed on a high-fat/fructose/cholesterol Western Diet (KT324, CorVus Biomedical, LLC) containing 16.3% kCal from protein, 42.9% kCal from fat, and 40.8% kCal from carbohydrates at approximately 2 months of age. Animals were fed once per day, and water was supplied ad libitum. All animal protocols were approved by the University of Missouri Animal Care and Use Committee and were in accordance with the "Principles for the Utilization of Care of Vertebrate Animals Used in Testing Research and Training".

**Ischemia/Reperfusion Induced Myocardial Infarction.** Ischemia/reperfusion induced myocardial infarction was performed as previously described with modifications [1,2]. At 6 months of age, pigs were sedated with 7.0 mg/kg Telazol intramuscular (IM), followed by 0.03 mg/kg buprenorphine IM for pain management. Anesthesia was induced with 2.0 mg/kg propofol intravenous (IV) and maintained by constant rate infusion (CRI) of 4-20 mg/kg/min. Pigs were mechanically ventilated with 100% oxygen at a rate of 8-12 breaths per minute and a tidal volume of 10-20 mL/kg and 20-25 cm $H_2O$ pressure. Following percutaneous femoral artery access, a 6F guide catheter (Boston Scientific) was advanced to the left main artery, and an appropriately sized angioplasty balloon (Abbott Trek) was placed into the left anterior descending coronary artery (LAD). The LAD was occluded distal to the 1$^{st}$ diagonal branch (D1) for 90 minutes, followed by reperfusion. During the ischemia-reperfusion procedure, ECG, MAP pressure, and other vital signs were continuously monitored, and cardioversion was utilized as necessary to prevent fatal arrythmia using a standard 200 Joule biphasic defibrillator.

**Echocardiography.** Transthoracic echocardiography was performed as previously described [1-5] pre- and immediately post-MI, 1-month post-MI, and 2-months post-MI using a GE Vivid I Ultrasound system with a 1.5 MHz transducer. Short-axis two-dimensional M-mode images were collected at the mid-papillary level of the LV and were analyzed offline using GE EchoPac Software.

**Osmotic-Pump Placement and FSTL1 delivery.** One month following MI, pigs were sedated with Telazol (7.0 mg/kg) IM, followed by 0.02 mg/kg buprenorphine IM for pain management. Anesthesia was maintained using isoflurane inhalation (~1.5-3%). Alzet Osmotic Pumps were implanted subcutaneously on both lateral surfaces of the neck. One Alzet pump released FSTL1 (120 mg FSTL1 in 2 ml TBS, released 5.0 uL per hour over 14 days) or vehicle (TBS, Tris Buffered Saline). The other Alzet pump released EdU (12 mg EdU in 2 ml PBS, released 2.5 uL per hour for 28 days) to label proliferating cells or vehicle (PBS, Phosphate Buffered Saline). Excede antibiotic (5.0 mg/kg) was given IM post-procedure.

**Pressure-Volume Loops.** Pressure-Volume (PV) loops were collected as previously described [2-6]. Two months following MI, pigs were sedated with a mixture of Telazol (5 mg/kg)/Xylazine (2.25 mg/kg), and anesthesia was maintained using propofol (6-14 mg/kg/min). Upon vascular access, heparin was given IV with an initial loading dose of 300 U/kg, followed by maintenance of 100 U/kg each hour. Following isolation and access of the femoral artery and vein, a 14F occlusion balloon catheter (Edward Life Sciences) was positioned in the inferior vena cava at the level of the apex of the heart, and a 6F guide catheter was positioned in the aorta for continuous monitoring of MAP. A median sternotomy was performed, and a small portion of the pericardium near the apex was opened for insertion of a calibrated 5F admittance-based Advantage pressure-volume (PV) loop catheter (Transonic Systems, Inc.; Ithaca, NY) into the LV. PV loops were recorded at rest and during conditions of reducing pre-load by transient occlusion of the inferior vena cava. LV pressure and LV volume were recorded as a function of time. LV PV loops were analyzed for end systolic and end diastolic pressures, volumes, and pressure volume relationships (ESPVR and EDPVR respectively), Preload Recruitable Stroke Work (PRSW), and Ventricular-Arterial Coupling Ratio [(end systolic pressure/stroke volume)/ESPVR].

**Microspheres.** Regional blood flow was measured by injecting 13-20 x$10^6$ Samarium-labeled microspheres (15μm diameter) (BioPAL,Inc., Worcester MA) into the left ventricular apex. An arterial reference sample was collected from the distal portion of the abdominal aorta at a withdrawal rate of 3 ml/min beginning prior to the injection of microspheres and continuing for a total of 3 minutes. After euthanasia, tissues were dissected from the animal, weighed, and dried in BioPal sample vials. Tissue and reference blood samples were sent to BioPAL and analyzed using neutron activation technology. Blood flow per 100 grams of tissue ($Q_m$) was calculated as: $Q_m = Q_r \times D_m/D_r$, where $Q_r$ is the rate of withdrawal of the reference blood sample (in ml/min), $D_m$ is the disintegrations per minute per gram of tissue, and $D_r$ is the disintegrations per minute of the reference blood sample.

**Pressure Myography.** Coronary arterioles from the infarct, border, and remote regions of the heart, as well as second order (2A) pial cerebral arterioles of the middle cerebral artery were isolated, cannulated, and pressurized for pressure myography as performed previously [7,8]. Constrictive capacity was assessed using U46619 (thromboxane A2 agonist), and dilatory capacity was assessed using NS1619 (large-conductance calcium-activated potassium channel; $BK_{Ca}$ activator).

**Mitochondrial Function.** Mitochondrial isolation and functional analysis were performed as previously described [9]. Portions of infarcted LV were placed in mitochondrial isolation buffer, homogenized with a Teflon pestle, and isolated using a percoll gradient and centrifugation. Mitochondrial respiration was assessed using high-resolution respirometry (Oroboros Oxygraph-2k; Oroboros Instruments). Basal respiration was assessed by loading mitochondria then adding substrates as previously described [9,10]. State 2 respiration was stimulated by the addition of malate (2 mM) and glutamate (5 mM), State 3 (maximal Complex I) respiration by titrated ADP (250-2000 uM), State 3 (Complex I & II) by addition of succinate (10 mM), and maximal coupling by addition of carbonyl cyanide-p-trifluoromethoxy phenylhydrazone (FCCP, 0.25 uM). Mitochondrial respiration was normalized to protein concentration obtained from BCA assay per manufacturer's instructions.

**Immunohistochemistry.** Immunohistochemistry was performed as previously described with modifications [11,12]. Formalin-fixed, paraffin embedded, 8um sections of left ventricle were stained with Hematoxylin & Eosin and Mallory's Prussian Blue Iron Stain.

**Immunofluorescence.** Immunofluorescence was performed as previously described [13]. Formalin-fixed, paraffin embedded, 8um sections of left ventricle were stained using ClickIt EdU, RBM20, and DAPI.

**Statistical Analyses.** All data were graphed and analyzed using GraphPad Prism 10. Pressure myograpy data, mitochondrial data, and serial echocardiography data were analyzed using repeated measures ANOVA, All other two group comparisons (MI vs MI + FSTL1) were analyzed using unpaired t-tests, Outliers were determined using ROUT with Q=5%. Data were considered significant at *p<0.05 and †p<0.1.

### References


1 Ishikawa, K. *et al.* Characterizing preclinical models of ischemic heart failure: differences between LAD and LCx infarctions. *Am J Physiol Heart Circ Physiol* **307**, H1478-1486, doi:10.1152/ajpheart.00797.2013 (2014).

2 Zaid, M. *et al.* Mechanism-Driven Modeling to Aid Non-invasive Monitoring of Cardiac Function via Ballistocardiography. *Front Med Technol* **4**, 788264, doi:10.3389/fmedt.2022.788264 (2022).

3 Olver, T. D. *et al.* Western Diet-Fed, Aortic-Banded Ossabaw Swine: A Preclinical Model of Cardio-Metabolic Heart Failure. *JACC Basic Transl Sci* **4**, 404-421, doi:10.1016/j.jacbts.2019.02.004 (2019).

4 Chade, A. R. *et al.* A New Model of Chronic Kidney Disease, Metabolic Derangements, and Heart Failure with Preserved Ejection Fraction in Aging Swine. *Am J Nephrol* **56**, 337-350, doi:10.1159/000543327 (2025).

5 Chade, A. R. *et al.* Chronic kidney disease and left ventricular diastolic dysfunction (CKD-LVDD) alter cardiac expression of mitochondria-related genes in swine. *Transl Res* **267**, 67-78, doi:10.1016/j.trsl.2023.12.004 (2024).

6 Olver, T. D. *et al.* Chronic interval exercise training prevents BK(Ca) channel-mediated coronary vascular dysfunction in aortic-banded miniswine. *J Appl Physiol (1985)* **125**, 86-96, doi:10.1152/japplphysiol.01138.2017 (2018).

7 Olver, T. D. *et al.* Loss of Female Sex Hormones Exacerbates Cerebrovascular and Cognitive Dysfunction in Aortic Banded Miniswine Through a Neuropeptide Y-Ca(2+)-Activated Potassium Channel-Nitric Oxide Mediated Mechanism. *J Am Heart Assoc* **6**, doi:10.1161/JAHA.117.007409 (2017).

8 Heaps, C. L., Tharp, D. L. & Bowles, D. K. Hypercholesterolemia abolishes voltage-dependent K+ channel contribution to adenosine-mediated relaxation in porcine coronary arterioles. *Am J Physiol Heart Circ Physiol* **288**, H568-576, doi:10.1152/ajpheart.00157.2004 (2005).

9 Sheldon, R. D. *et al.* eNOS deletion impairs mitochondrial quality control and exacerbates Western diet-induced NASH. *Am J Physiol Endocrinol Metab* **317**, E605-E616, doi:10.1152/ajpendo.00096.2019 (2019).

10 Kelty, T. J. *et al.* Western diet-induced obesity results in brain mitochondrial dysfunction in female Ossabaw swine. *Front Mol Neurosci* **16**, 1320879, doi:10.3389/fnmol.2023.1320879 (2023).

11 Tharp, D. L. *et al.* Local delivery of the KCa3.1 blocker, TRAM-34, prevents acute angioplasty-induced coronary smooth muscle phenotypic modulation and limits stenosis. *Arterioscler Thromb Vasc Biol* **28**, 1084-1089, doi:10.1161/ATVBAHA.107.155796 (2008).

12 Bowles, D. K., Heaps, C. L., Turk, J. R., Maddali, K. K. & Price, E. M. Hypercholesterolemia inhibits L-type calcium current in coronary macro-, not microcirculation. *J Appl Physiol (1985)* **96**, 2240-2248, doi:10.1152/japplphysiol.01229.2003 (2004).

13 Wei, K. *et al.* Epicardial FSTL1 reconstitution regenerates the adult mammalian heart. *Nature* **525**, 479-485, doi:10.1038/nature15372 (2015).
